# Functional brain network modeling in sub-acute stroke patients and healthy controls during rest and continuous attentive tracking

**DOI:** 10.1101/644765

**Authors:** Erlend S. Dørum, Tobias Kaufmann, Dag Alnæs, Geneviève Richard, Knut K. Kolskår, Andreas Engvig, Anne-Marthe Sanders, Kristine Ulrichsen, Hege Ihle-Hansen, Jan Egil Nordvik, Lars T. Westlye

**Author notes:** Corresponding authors: Erlend S. Dørum & Lars T. Westlye, Department of Psychology, University of Oslo, PoBox 1094 Blindern, 0317 OSLO, Norway.

## Abstract

A cerebral stroke is characterized by compromised brain function due to an interruption in cerebrovascular blood supply. Although stroke incurs focal damage determined by the vascular territory affected, clinical symptoms commonly involve multiple functions and cognitive faculties that are insufficiently explained by the focal damage alone. Functional connectivity (FC) refers to the synchronous activity between spatially remote brain regions organized in a network of interconnected brain regions. Functional magnetic resonance imaging (fMRI) has advanced this system-level understanding of brain function, elucidating the complexity of stroke outcomes, as well as providing information useful for prognostic and rehabilitation purposes.

We tested for differences in brain network connectivity between a group of patients with minor ischemic strokes in sub-acute phase (n=44) and matched controls (n=100). As neural network configuration is dependent on cognitive effort, we obtained fMRI data during rest and two load levels of a multiple object tacking (MOT) task. Network nodes and time-series were estimated using independent component analysis (ICA) and dual regression, with network edges defined as the partial temporal correlations between node pairs. The full set of edgewise FC went into a cross-validated regularized linear discriminant analysis (rLDA) to classify groups and cognitive load.

MOT task performance and cognitive tests revealed no significant group differences. While multivariate machine learning revealed high sensitivity to experimental condition, with classification accuracies between rest and attentive tracking approaching 100%, group classification was at chance level, with negligible differences between conditions. Repeated measures ANOVA showed significantly stronger synchronization between a temporal node and a sensorimotor node in patients across conditions. Overall, the results revealed high sensitivity of FC indices to task conditions, and suggest relatively small brain network-level disturbances after clinically mild strokes.

## Introduction

Unlike the insidious onset and progressive neurological decline observed in most neurodegenerative diseases, a cerebral stroke is characterized by instant damage to brain tissue due to a compromise in cerebrovascular blood supply. Although stroke incurs focal damage determined by the vascular territory affected, clinical symptoms commonly involve multiple functions and cognitive faculties that are insufficiently explained by the focal damage alone (Carter et al., 2012; Ovadia-Caro et al., 2013).

The brain is organized in a network of connected brain regions that are spatially dispersed, yet functionally linked (Bullmore & Sporns, 2009; Power et al., 2011; Van Den Heuvel & Pol, 2010), thus even a well localized stroke can give rise to a complex clinical picture of symptoms owing to the highly interconnected organization of the cerebral cortex. Contemporary brain network connectivity models go beyond traditional lesion-symptom mapping, and advanced imaging techniques like functional magnetic resonance imaging (fMRI) can be used to capture disruptions of connectivity following stroke. fMRI has shown great promise in probing alterations in brain activity for a range of neurodegenerative (Cordova-Palomera et al., 2017; Dickerson et al., 2016; Paulsen et al., 2004) and neuropsychiatric (Kaufmann et al., 2015; Skåtun et al., 2016; Yu et al., 2015) conditions. These techniques may provide novel understanding of brain function, and potentially, clinical information used to predict patient recovery, outcome and aid in tailoring individual rehabilitation strategies.

Functional connectivity (FC) refers to the temporal correlation of blood-oxygen-level dependent (BOLD) signal between brain regions (Hampson et al., 2002). Resting state fMRI has revealed the brain to be organized into functional networks of distributed brain systems, where an orchestra of synchronous activity underlies even the simplest behaviors (Van Den Heuvel & Pol, 2010). Task based fMRI studies have shown that functional connections at rest are similarly engaged during various cognitive tasks (Raichle, 2010; Smith et al., 2009) and the increase in joint BOLD activations in spatially dispersed brain regions has revealed networks engaged during attention (Alnæs et al., 2015; Szczepanski et al., 2013), working memory (Compte et al., 2000), language (Ferstl et al., 2008) and motor task (Hanakawa et al., 2008), as well as various other cognitive operations.

Most studies relating the effect of stroke on fMRI-based FC to behavioral deficits have used unconstrained resting state data. Siegel et al. (2016) showed that FC was superior in predicting certain memory deficits, whilst visual and motor impairments were best predicted by lesion topography. Attention and language deficits were well predicted by both. He and colleagues (2007) demonstrated that the severity of spatial neglect was correlated with the degree of disruption within the contralateral attention network connectivity.

The present work builds on a previous study that examined age differences in functional connectivity in healthy controls during 1) an unconstrained resting-state condition and 2) two load levels of a constrained continuous tracking task (Dørum et al., 2017). Briefly, our previous findings demonstrated that a machine learning approach based on FC resulted in robust discrimination between a group of younger and a group of older healthy participants, as well as between states of rest and effortful attention, with higher sensitivity to age group observed during constrained tracking compared to resting state. In the present study, we aim to apply a similar prediction model on data from 44 patients with stroke and 100 healthy controls, in order to test whether the functional brain data alone would be sufficient to accurately identify the stroke patients, and whether the group classification varied between load levels. We also investigated edge-level main effects of experimental condition and group, as well as their interactions, using repeated measures ANOVA. Lastly, since edge-level FC ultimately reflects nodal signal changes, temporal activity on node-level was probed by computing the standard deviation of signal amplitude (SDSA) (Garrett et al., 2010). Brain signal variability reflects neural complexity and the ability to dynamically transition between a range of network states (Garrett et al., 2011), and a reduction in signal variability has been reported with advancing age (Grady & Garrett, 2014), in patients with schizophrenia (Kaufmann et al., 2015), and in acute stroke patients (Zappasodi et al., 2014), indicating global loss of complexity and neurological dysfunction.

Based on the studies reviewed above and current models of brain network dysfunction after stroke, we hypothesized 1) that multivariate classification would yield robust discrimination between a group of sub-acute stroke patients and healthy controls, as well as between states of rest and task engagement. Next, we anticipated that 2) FC during task would yield higher classification accuracy when classifying between stroke patients and healthy controls. We accompanied the multivariate analyses with edgewise repeated measures ANOVAs to test for effects of group, task condition (rest, L1, L2) and their interactions. Lastly, we hypothesized 3) decreased SDSA for stroke patients compared to healthy controls, with group differences more prominent during task compared to rest.

## Methods and material

### General study design

In this cross-sectional study, stroke patients and healthy controls underwent MRI examination, including a resting state and task-based functional image acquisition as well as cognitive and neuropsychological assessment.

### Study material and recruitment procedures

Patients were recruited from the stroke units at Oslo University hospital (OUS), Diakonhjemmet hospital and Bærum hospital, Norway. Inclusion criteria were: (i) age 18 or older, (ii) clinically and radiologically documented stroke of ischemic, hemorrhagic or subarachnoid origin, (iii) time of enrolment within 14 days of admittance. Exclusion criteria were: (i) clinical condition of impaired consciousness leading to inability to actively participate and maintain wakefulness psychiatric condition (e.g., schizophrenia, bipolar disorder) as well as alcohol or substance abuse that potentially could impact the interpretation of the behavioral/imaging data, (iii) contraindication for MRI including incompatible metal implants, claustrophobia or pregnancy.

Clinical assessment quantifying stroke severity was performed according to the National Institute of Health Stroke Scale (NIHSS) at the respective stroke units by an attending physician specialized in internal medicine, neurology or geriatric medicine at the time of discharge. Cognition was assessed using the Montreal Cognitive Assesment (MoCA) test after the patients were clinically stable and before discharge. The ischemic strokes were classifyed according to the TOAST classification (Adams et al., 1993). The patients were treated in accordance with national guidelines (Helsedirektoratet, 2017).

Healthy controls were recruited from social media and newspaper ads. Inclusion criteria were: (i) age 18 years or older, (ii) absence of neurologic, psychiatric condition as well as alcohol or substance abuse, (iii) abnormal radiological findings requiring medical follow-up (e.g. silent stroke, tumor). From a pool of 341 healthy controls recruited to a parallel study (Richard et al., 2018), we selected n=100 adults based on a group-level matching with the patient sample with regards to age, sex, education and handedness.

Written informed consent was obtained from all participants, and the Regional Committees for Medical Research Ethics South East Norway approved the study protocol.

### MRI acquisition

MRI scans were obtained using a General Electric Medical Systems (Discovery MR750) 3.0T scanner with a 32-channel head coil at Oslo University Hospital. fMRI data was acquired with a T2*-weighted 2D gradient echo planar imaging sequence (EPI) (TR: 2250 ms; TE: 30 ms; FA: 79°; voxel size: 2.67 × 2.67 ×3.0 mm; slices: 43; FOV: 96 × 96 × 129 mm. We collected 200 volumes for the resting-state condition and 152 volumes for the two MOT load conditions, after discarding the first five volumes. We collected a structural scan using a sagittal T1-weighted fast spoiled gradient echo (FSPGR) sequence (TR: 8.16 ms; TE: 3.18 ms; TI: 450 ms; FA: 12°; voxel size: 1.0 ×1.0×1.0 mm; slices: 188; FOV: 256 × 256 × 188 mm; duration: 288 s), and a T2-FLAIR (TR: 8000 ms; TE: 127 ms, TI: 2240; voxel size: 1.0 × 1.0 × 1.0 mm; duration 443 s) for lesion demarcation.

### fMRI paradigms

Participants underwent one resting state run and three versions of MOT, including one blocked and two continuous tracking runs, performed in the MRI scanner during the same session (Alnæs et al., 2015). Here we report results from the resting-state and the two continuous load conditions. The level of attentional demand was set at two load conditions – load 1 (L1) and load 2 (L2) requiring the participants to track one or two targets, respectively. We restricted the load level to 2 to ensure that both groups were able to perform at a high level.

The task was presented on a calibrated MR-compatible LCD screen (NNL LCD Monitor®, NordicNeuroLab, Bergen, Norway) with a screen resolution of 1920×1080 at 60Hz, placed in front of the scanner bore. The experimental set-up and technical specifications were performed as described in a previous publication (Alnæs et al., 2015).

Each participant performed two runs of continuous MOT task, one with tracking load of 1 object, and the other with 2 objects. Each continuous tracking block lasted 7.5 minutes and contained 14 trials. Detailed outline of the task is described in our previous study (Dørum et al., 2017). Briefly, 10 identical blue objects were presented on a grey background screen, and after another 0.5 seconds the objects started moving. The first cued object(s), or target(s), turned red after the first 0.5 second, and remained red for a duration of 2.5 seconds. The tracking period lasted on average 32 seconds (range, 27-37) after which the participants were instructed to respond to a probe (green object), “yes” or “no” to whether the green probe was one of the objects originally designated as a target, before a new target assignment took place. The participants were instructed to fixate on a central fixation point during the length of the run. Accuracy and reaction times were recorded.

### Lesion demarcation

Individual lesions were defined based on visible damage and hyperintensities on FLAIR images as well as guided by independent neuroradiological descriptions using DWI/FLAIR images. The lesions were semi-automatically delineated in native space using the Clusterize toolbox (de Haan et al., 2015) used with SPM8, running under Matlab R2013b (The Mathworks, Inc., Natick, MA). The FLAIR images were registered with the high-resolution T1 images using a linear transformation with 6 degrees-of-freedom. Subsequently, each T1 image was registered to the MNI152 standard space by computing 12 degrees-of-freedom linear affine transformation. To obtain each registered lesion mask in standard space, the native-to-standard transformation matrices were applied using the nearest neighbor interpolation. Figure 1 shows a probabilistic lesion heat-map displaying lesion overlap across patients.

**Figure 1.**
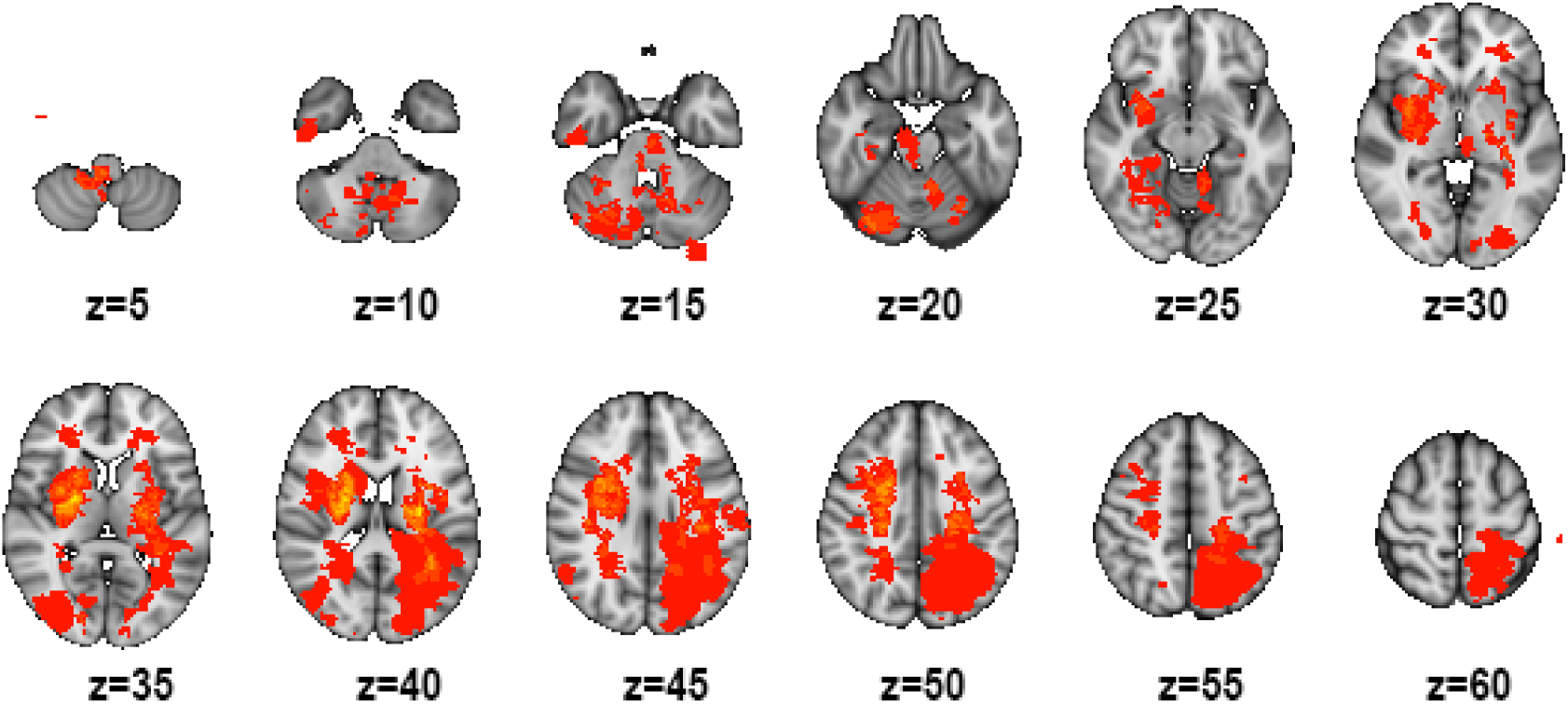
Probabilistic lesion heat-map across all stroke patients with coordinates in MNI-space. Colors towards the yellow range indicate higher degree of lesion overlap across the stroke group. Left-right orientation is in accordance to radiological convention.

### fMRI analysis

FMRI data was processed on single-subject level using the FMRI Expert Analysis Tool (FEAT) from the FMRIB Software Library (FSL) (Smith et al., 2004) including spatial smoothing (FWHM = 6 mm), high-pass filtering (sigma = 64 s), motion correction (MCFLIRT) and single-session independent component analysis (ICA) using MELODIC (Beckmann & Smith, 2004). In-scanner motion was calculated as the average root mean square of the displacement from one frame to its previous frame for each dataset. Independent samples t-tests revealed no significant difference in motion during rest [mean(SD)_controls_ = .09 (.05), mean(SD)_patients_ = .11 (.08), t = −1.54 p = .13]; during L1 [mean(SD)_controls_ = .11 (.08), mean(SD)_patients_ = .13 (.09), t = −1.50, p = .14]; and during L2 [mean(SD)_controls_ = .14 (.14), mean(SD)_patients_ = .15 (.09), t = -.65, p = .52]

We used FSL FIX (FMRIB’s ICA-based Xnoisefier) (Salimi-Khorshidi et al., 2014) to identify and remove noise components at the individual level (standard training set, threshold: 20), and regressed out the estimated motion parameters from MCFLIRT from the voxel-wise time series.

Next, we employed a group-level PCA approach (Smith et al., 2014) in MELODIC to estimate group-level spatial maps representing the nodes in our networks. We used a model order of 40 and discarded 10 noise components based on the spatial distribution of the component maps and/or the frequency spectrum of the components’ time series (Kelly et al., 2010), resulting in a final set of 30 components (see Fig 2). Next, the full set of spatial maps from the group analysis was used to generate subject-specific versions of the spatial maps, and associated timeseries, using dual regression (Nickerson et al., 2017). First, for each subject, the group-average set of spatial maps is regressed (as spatial regressors in a multiple regression) into the subject’s 4D space-time dataset. This results in a set of subject-specific timeseries, one per group-level spatial map. Next, those timeseries are regressed (as temporal regressors, again in a multiple regression) into the same 4D dataset, resulting in a set of subject-specific spatial maps, one per group-level spatial map. Here, in order to remove common variance with the lesion, we included the segmented lesion mask as an additional component in individual dual regression runs.

**Figure 2:**
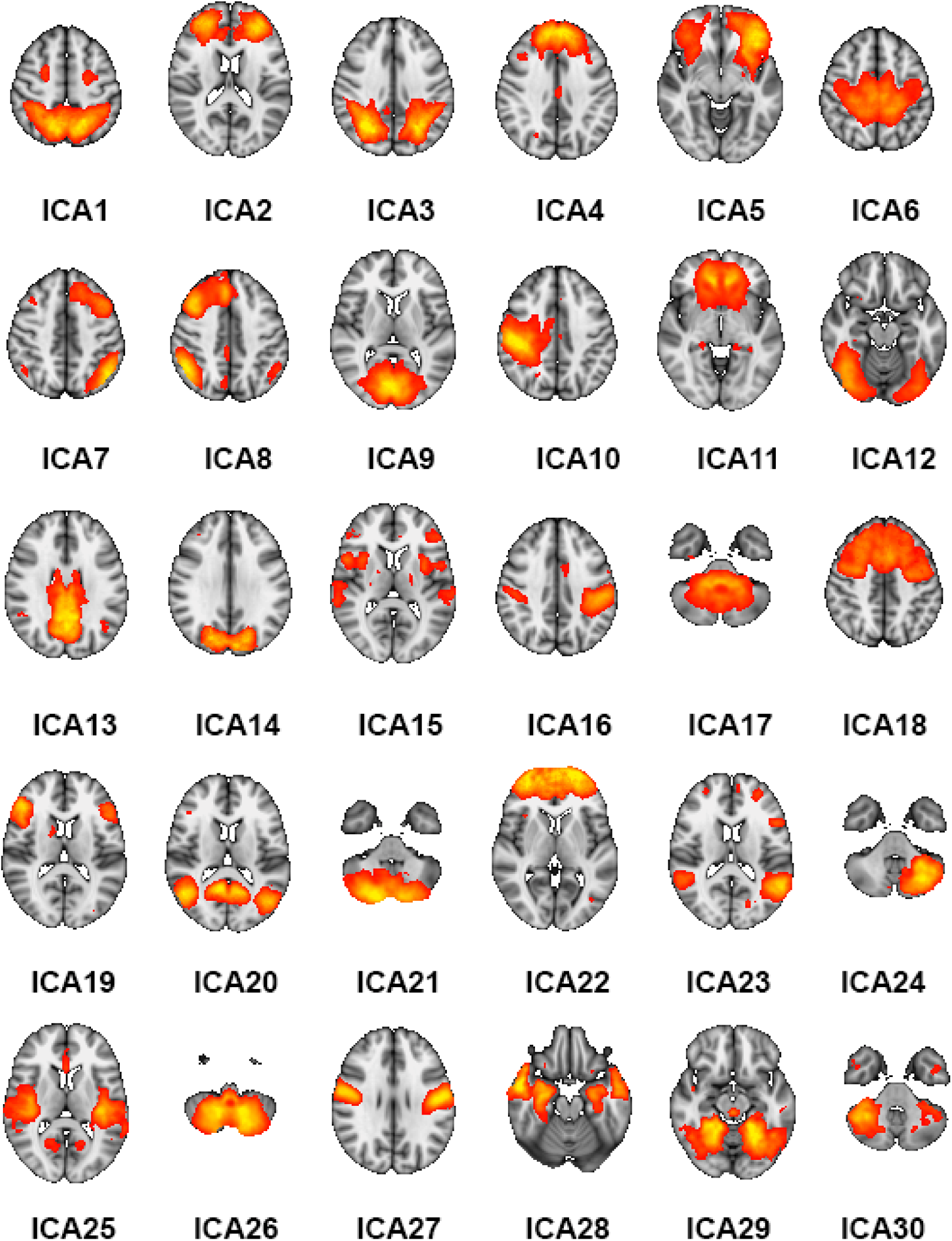
The 30 networks derived from the ICA numbered from top left to bottom right.

After discarding the estimated time-series from the lesion, we used the components’ time series for network modeling (Smith et al., 2011). Here, the spatial maps are considered nodes in an extended brain network, and the edges are defined as the temporal correlation between each pair of nodes. Based on our previous work (Kaufmann et al., 2016), we estimated the temporal correlations using regularized partial correlations with an automated lambda estimation (Brier et al., 2015; Ledoit & Wolf, 2003). This approach resulted in 435 unique edges for each individual, which were submitted to further group-level univariate and multivariate analyses (see below).

For each dataset, we also computed the individual level SDSA of each component’s time series as a measure of nodal volatility or strength (Garrett et al., 2010; Kaufmann et al., 2015).

### Statistical analysis

Demographics, behavioral and clinical data were analyzed in SPSS (IBM_Corp, 2010). Between-group differences were assessed using Chi square tests (sex distribution) and linear models (age, neuropsychological performance, MOT task performance).

In order to assess the predictive value of the FC measures on the individual level we employed a multivariate machine learning approach based on our previous work (Alnæs et al., 2015; Dørum et al., 2017). We used the network edges to classify task condition across and within groups, and also classified group (case/control) across and within task conditions, using a regularized linear discriminant classifier (shrinkage LDA) (Friedman, 1989; Schäfer & Strimmer, 2005). To avoid bias due to the uneven group sizes, we performed the group classification within a nested loop of 100 iterations, in which we each time randomly picked healthy controls to match the sample size of the patient group. The robustness of the models was assessed using leave-one-out cross-validation to avoid overfitting and permutation testing across 10,000 iterations to compare the predictive value to empirical null distributions.

In order to assess the univariate associations at the edge level, we performed repeated measures ANOVA to test for effects of group, task condition (rest, L1, L2) for each edge, and their interactions. We computed edgewise F-stats and adjusted the alpha level using false discovery rate (FDR) for each test separately with an FDR level q=0.05 and a threshold based on the assumption of independence or positive dependence (Nichols, 2009; Nichols & Hayasaka, 2003).

## Results

### Sample descriptives

We included 44 patients with ischemic stroke in the present analyses. Of the 54 patients initially enrolled in the study, 10 were excluded (3 were diagnosed with diseases other than stroke, 3 had stroke-related visual and/or motor impairments rendering them incapable of performing the MOT task, 4 were unable to complete the MRI protocol and/or the neuropsychological tests). Table 1 provides individual patient level information regarding lesion location, stroke classification and days between stroke incident and MRI scan.

**Table 1.**
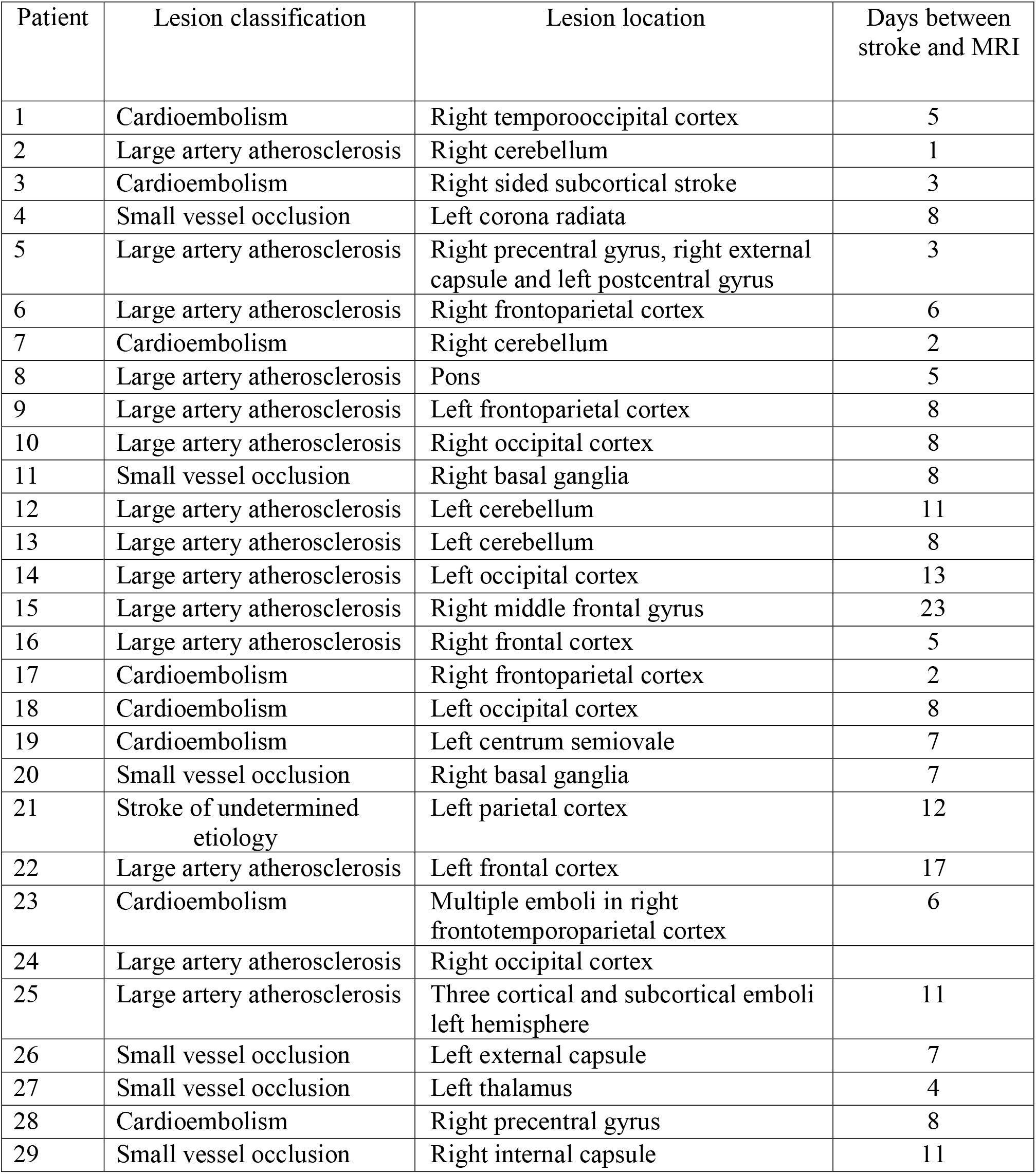
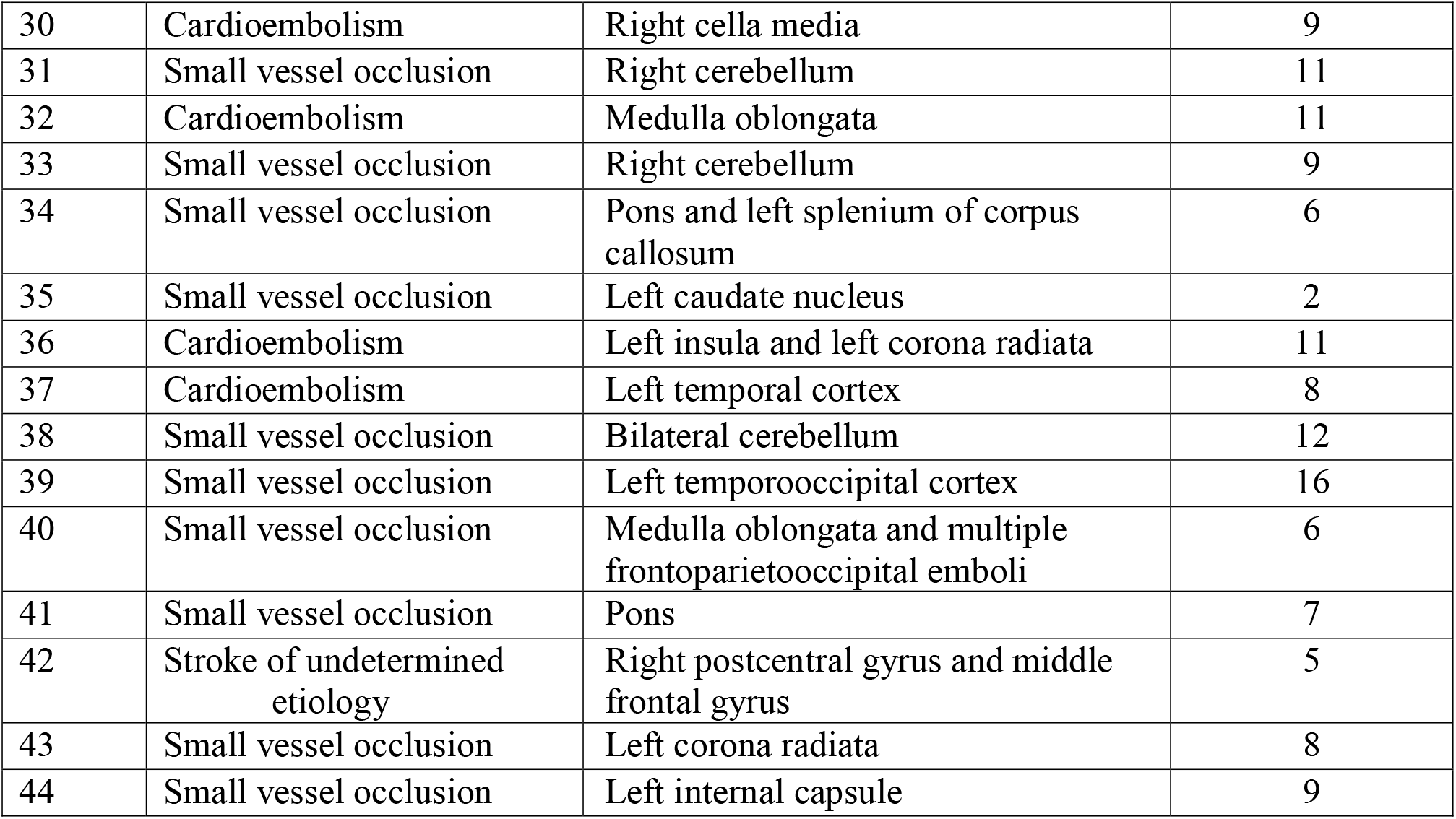
Patient sample with classification according to TOAST, lesion location and days between stroke incident and MRI scan.

| Patient | Lesion classification | Lesion location | Days between stroke and MRI |
| --- | --- | --- | --- |
| 1 | Cardioembolism | Right temporooccipital cortex | 5 |
| 2 | Large artery atherosclerosis | Right cerebellum | 1 |
| 3 | Cardioembolism | Right sided subcortical stroke | 3 |
| 4 | Small vessel occlusion | Left corona radiata | 8 |
| 5 | Large artery atherosclerosis | Right precentral gyrus, right external capsule and left postcentral gyrus | 3 |
| 6 | Large artery atherosclerosis | Right frontoparietal cortex | 6 |
| 7 | Cardioembolism | Right cerebellum | 2 |
| 8 | Large artery atherosclerosis | Pons | 5 |
| 9 | Large artery atherosclerosis | Left frontoparietal cortex | 8 |
| 10 | Large artery atherosclerosis | Right occipital cortex | 8 |
| 11 | Small vessel occlusion | Right basal ganglia | 8 |
| 12 | Large artery atherosclerosis | Left cerebellum | 11 |
| 13 | Large artery atherosclerosis | Left cerebellum | 8 |
| 14 | Large artery atherosclerosis | Left occipital cortex | 13 |
| 15 | Large artery atherosclerosis | Right middle frontal gyrus | 23 |
| 16 | Large artery atherosclerosis | Right frontal cortex | 5 |
| 17 | Cardioembolism | Right frontoparietal cortex | 2 |
| 18 | Cardioembolism | Left occipital cortex | 8 |
| 19 | Cardioembolism | Left centrum semiovale | 7 |
| 20 | Small vessel occlusion | Right basal ganglia | 7 |
| 21 | Stroke of undetermined etiology | Left parietal cortex | 12 |
| 22 | Large artery atherosclerosis | Left frontal cortex | 17 |
| 23 | Cardioembolism | Multiple emboli in right frontotemporoparietal cortex | 6 |
| 24 | Large artery atherosclerosis | Right occipital cortex |  |
| 25 | Large artery atherosclerosis | Three cortical and subcortical emboli left hemisphere | 11 |
| 26 | Small vessel occlusion | Left external capsule | 7 |
| 27 | Small vessel occlusion | Left thalamus | 4 |
| 28 | Cardioembolism | Right precentral gyrus | 8 |
| 29 | Small vessel occlusion | Right internal capsule | 11 |
| 30 | Cardioembolism | Right cella media | 9 |
| 31 | Small vessel occlusion | Right cerebellum | 11 |
| 32 | Cardioembolism | Medulla oblongata | 11 |
| 33 | Small vessel occlusion | Right cerebellum | 9 |
| 34 | Small vessel occlusion | Pons and left splenium of corpus callosum | 6 |
| 35 | Small vessel occlusion | Left caudate nucleus | 2 |
| 36 | Cardioembolism | Left insula and left corona radiata | 11 |
| 37 | Cardioembolism | Left temporal cortex | 8 |
| 38 | Small vessel occlusion | Bilateral cerebellum | 12 |
| 39 | Small vessel occlusion | Left temporooccipital cortex | 16 |
| 40 | Small vessel occlusion | Medulla oblongata and multiple frontoparietooccipital emboli | 6 |
| 41 | Small vessel occlusion | Pons | 7 |
| 42 | Stroke of undetermined etiology | Right postcentral gyrus and middle frontal gyrus | 5 |
| 43 | Small vessel occlusion | Left corona radiata | 8 |
| 44 | Small vessel occlusion | Left internal capsule | 9 |

Table 2 summarizes key demographics, neuropsychological data and MOT performance for both groups, as well as stroke severity for the patient group as assessed by NIHSS. There were no significant group differences in any demographic variables, MoCA scores or MOT performance between patients and healthy controls.

**Table 2.** Demographics, stroke severity, and neuropsychologic performance. Standard deviations in parentheses.

| | Stroke | Healthy controls | $X^2 / t$ | p |
| --- | --- | --- | --- | --- |
| N | 44 | 100 |  |  |
| Age | 63.11 (14.8) | 63.12 (11.2) | $t = .00$ | .99 |
| Age range | 34 - 87 | 35 - 81 |  |  |
| Percent male | 75.0 | 60.0 | $X^2 = 3.00$ | .08 |
| Percent righthanded | 92.0 | 90.90 | $X^2 = .48$ | .83 |
| Years of education | 15.18 (2.0) | 15.82 (2.9) | $t = 1.52$ | .13 |
| MoCA | 26.91 (2.6) | 27.65 (1.7) | $t = 1.75$ | .09 |
| NIHSS | .73 (1.17) | NA |  |  |
| MOT accuracy L1 | 77.7 (24.9) | 84.0 (24.0) | $t = 1.42$ | .16 |
| MOT response time L1 | 1.13 (0.2) | 1.16 (0.2) | $t = .66$ | .51 |
| MOT accuracy L2 | 66.5 (23.9) | 71.7 (23.5) | $t = 1.22$ | .23 |
| MOT response time L2 | 1.17 (0.3) | 1.22 (0.2) | $t = 1.04$ | .30 |

### Classification analysis

Figure 3 shows the confusion matrices from all classification analyses. The algorithm distinguished resting state from both load conditions with high accuracy (mean classification accuracy across groups: 95.83%, p_perm_ < .0001, chance level = 30%). Classification of load conditions revealed minimal group differences, specifically L1 yielded slightly higher accuracy in the control group (L1: 67% p_perm_ < .0001) than in the stroke group (L1: 61.36%, p_perm_ < .0001), for load condition L2 classification accuracy was slightly higher for the stroke group (L2: 63.64%, p_perm_ < .0001) than the healthy control group (L2: 58.00%, p_perm_ < .0001).

**Figure 3:**
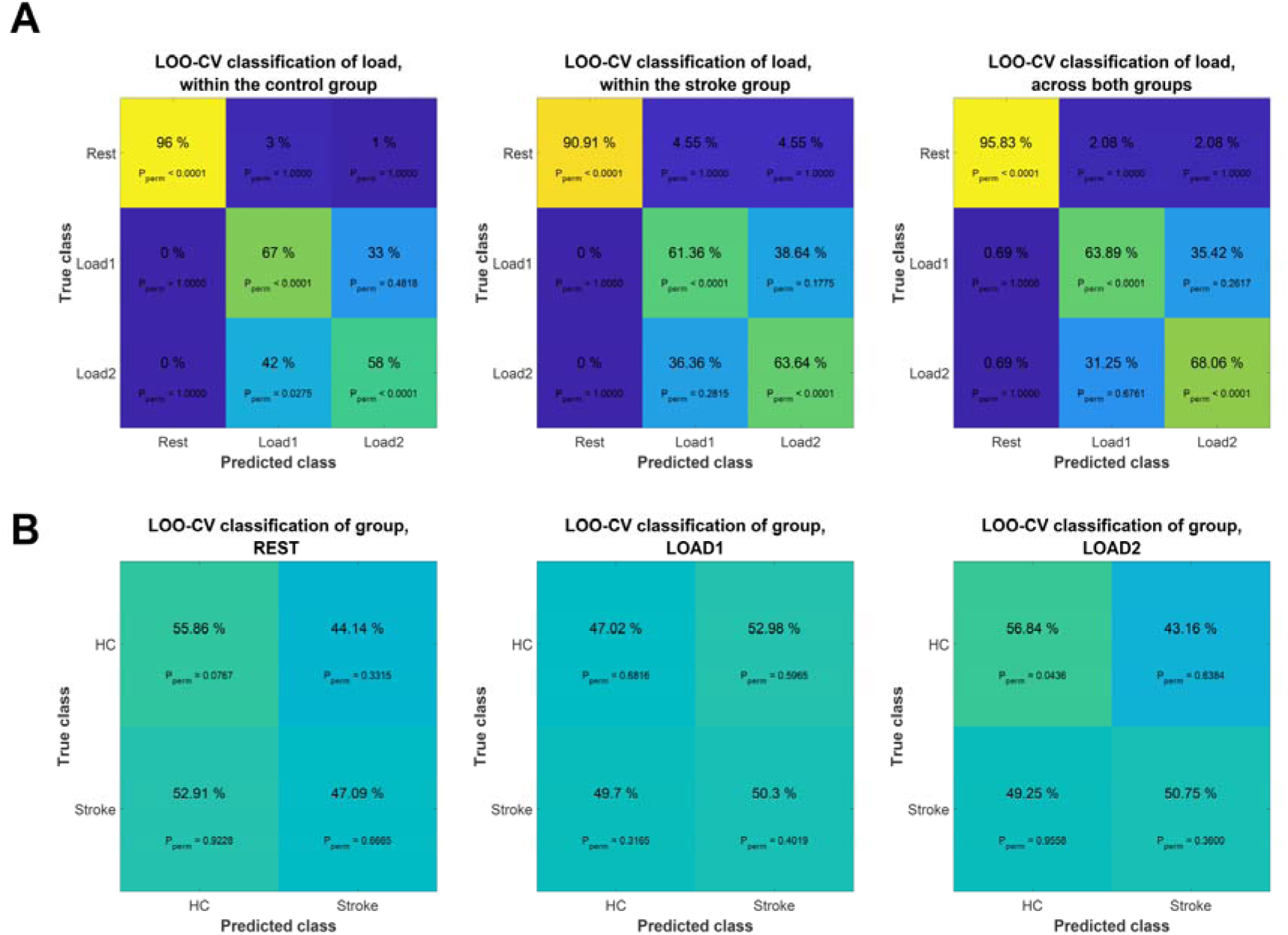
Confusion matrices from various classification tasks. A) Classification of the three conditions (rest, L1 and L2) within the stroke group, the healthy control group and across groups. B) Classification of the two groups (stroke and healthy controls) within resting state, load 1 and load 2.

We further trained the classifier to distinguish stroke patients from healthy controls based on data from resting state or each of the two load conditions separately. Briefly, classification performance was low, and, except for stroke group accuracy at L2, none of the classification tasks performed substantially better than chance level as estimated using permutation testing. Using resting state data, we achieved 47.09 % classification accuracy for identifying the stroke group (p_perm_ = .67) and 55.86% accuracy for identifying healthy controls (p_perm_ = .08). Accuracy when using L1 was 50.3% (p_perm_ = .40) and 47.02% (p_perm_ = .68) for the stroke and healthy control group, respectively. Accuracy using L2 was 50.75% (p_perm_ < .36) for patients and 56.84% (p_perm_ = .04) for controls.

### Edgewise univariate analysis

Figure 4 and Figure 5 summarize the results from the repeated measures ANOVA testing for A) main effect of condition, B) main effect of group and C) group by condition interaction effect. Repeated measures ANOVA revealed 210 edges showing significant main effect of condition. The edges showing strongest effect of condition were observed between nodes 14-20 (visual-DMN; F = 122.62, p < .001), nodes 1-7 (DAN-left frontoparietal; F = 119.53, p < .001), nodes 1-16 (DAN-supramarginal gyrus; F = 117.76, p < .001), nodes 1-27 (DAN-sensorimotor; F = 113.56, p < .001) and nodes 10-13 (right sensorimotor-DMN; F = 96.78, p < .001).

**Figure 4.**
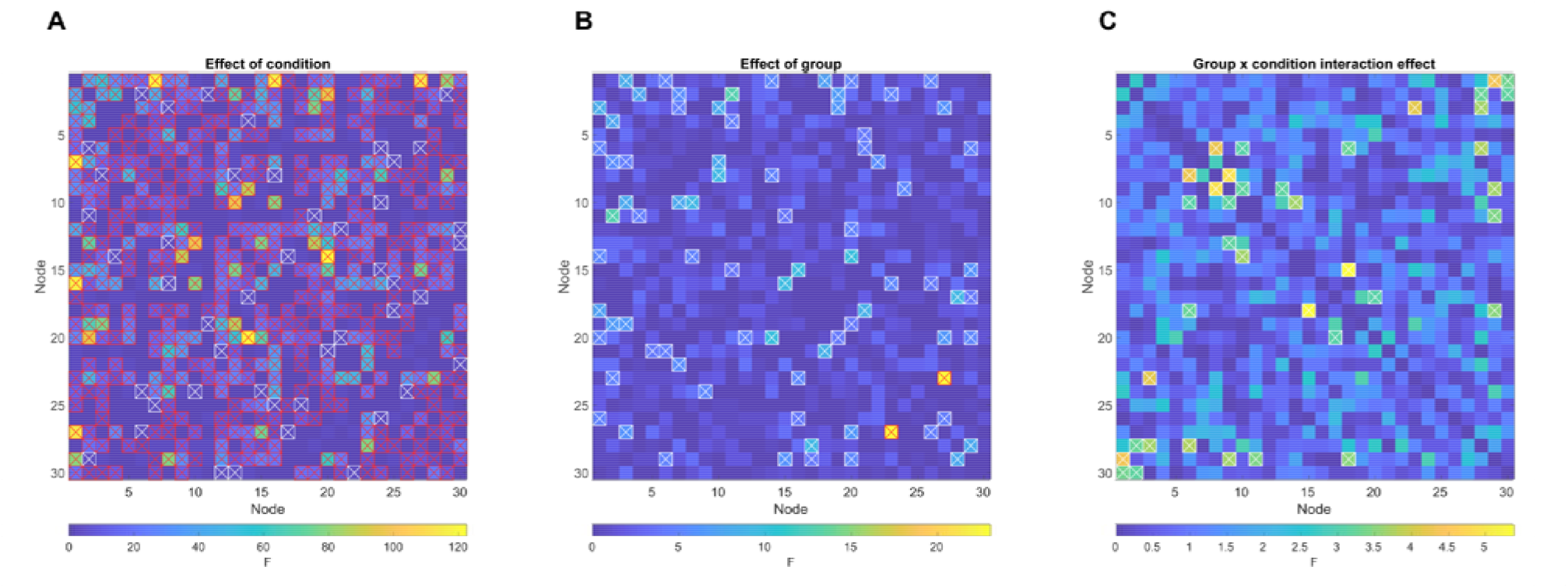
Edgewise repeated measures ANOVA. A) Main effect of condition, B) Main effect of group, C) Group by condition interaction effect. Red-crossed boxes denotes FDR-significant edges; white-crossed boxes denote nominally significant (p < .05) edges.

**Figure 5.**
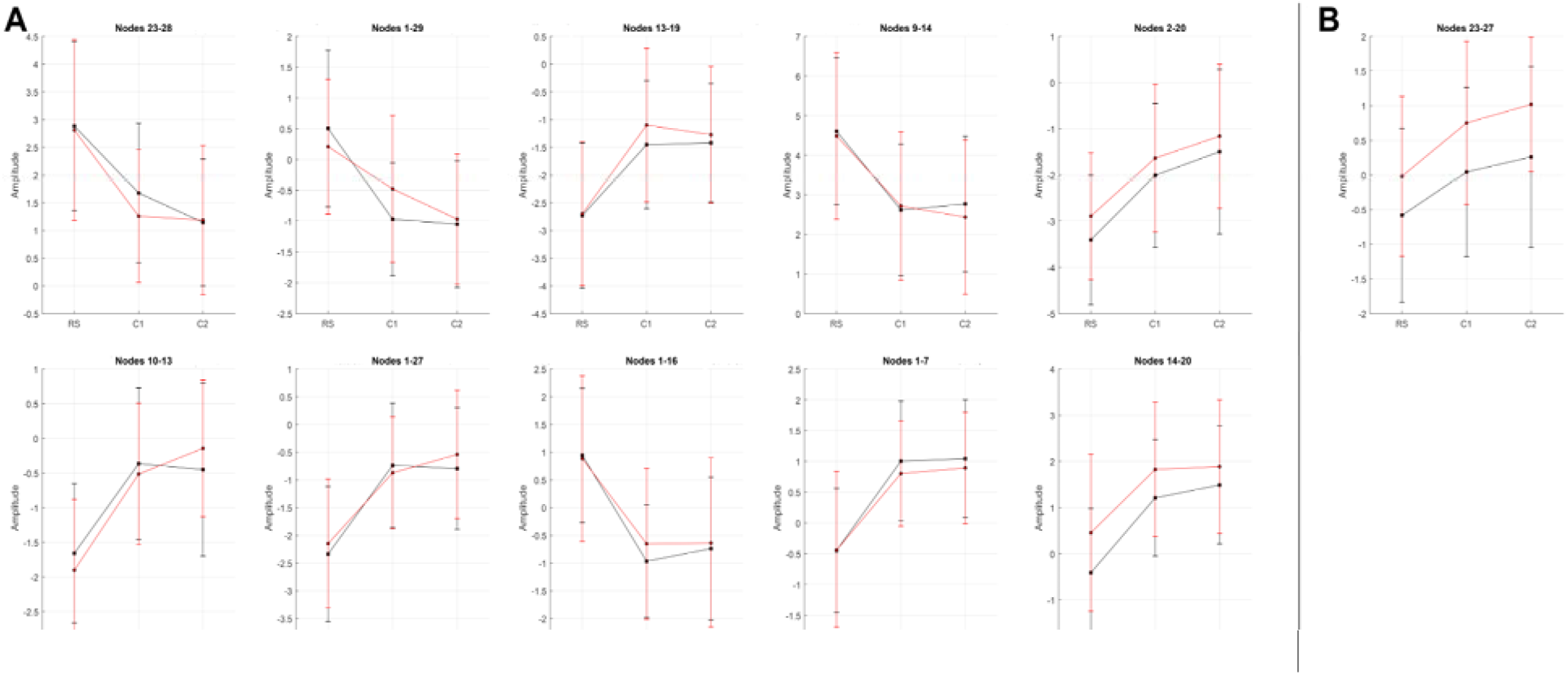
Changes in functional connectivity for stroke patients (red lines) and healthy controls (black lines) during the three conditions for A) the 10 edges showing strongest effect of condition and B) the single edge showing FDR-significant group effect

Among 41 edges showing nominally significant group differences, one edge remained significant after correction for multiple comparisons (p < .05, adjusted using FDR). The significant edge connected nodes 23 and 27 (temporal-sensorimotor; F = 23.08, p < .001), and showed a task-dependent increase in connectivity strength for both groups, with significantly higher connectivity in the stroke group during resting state and both task loads.

18 edges showed nominally significant group by condition interactions, however none survived FDR correction.

### SDSA

Figure 6 visualizes the results of the nodewise SDSA analysis. Repeated measures ANOVA revealed a significant (p < .05, FDR corrected) main effect of condition for 19 nodes. Strongest effect of condition was observed in node 20 [(inferior frontal gyrus), F = 205.96, p < .001], node 15 [(DAN), F = 150.00, p < .001], node 13 [(DMN), F = 104.98, p < .001], node 24 [(cerebellum), F = 57.85, p < .001] and node 23 [(temporal lobe), F = 48.64, p < .001]. Briefly, the frontal gyrus, DAN, DMN and temporal lobe nodes showed a task-related decrease in signal variability whereas a task-related increase was observed in the cerebellar node. FDR adjustment revealed no significant main effects of group and no significant interaction between group and condition on node-wise SDSA.

**Figure 6.**
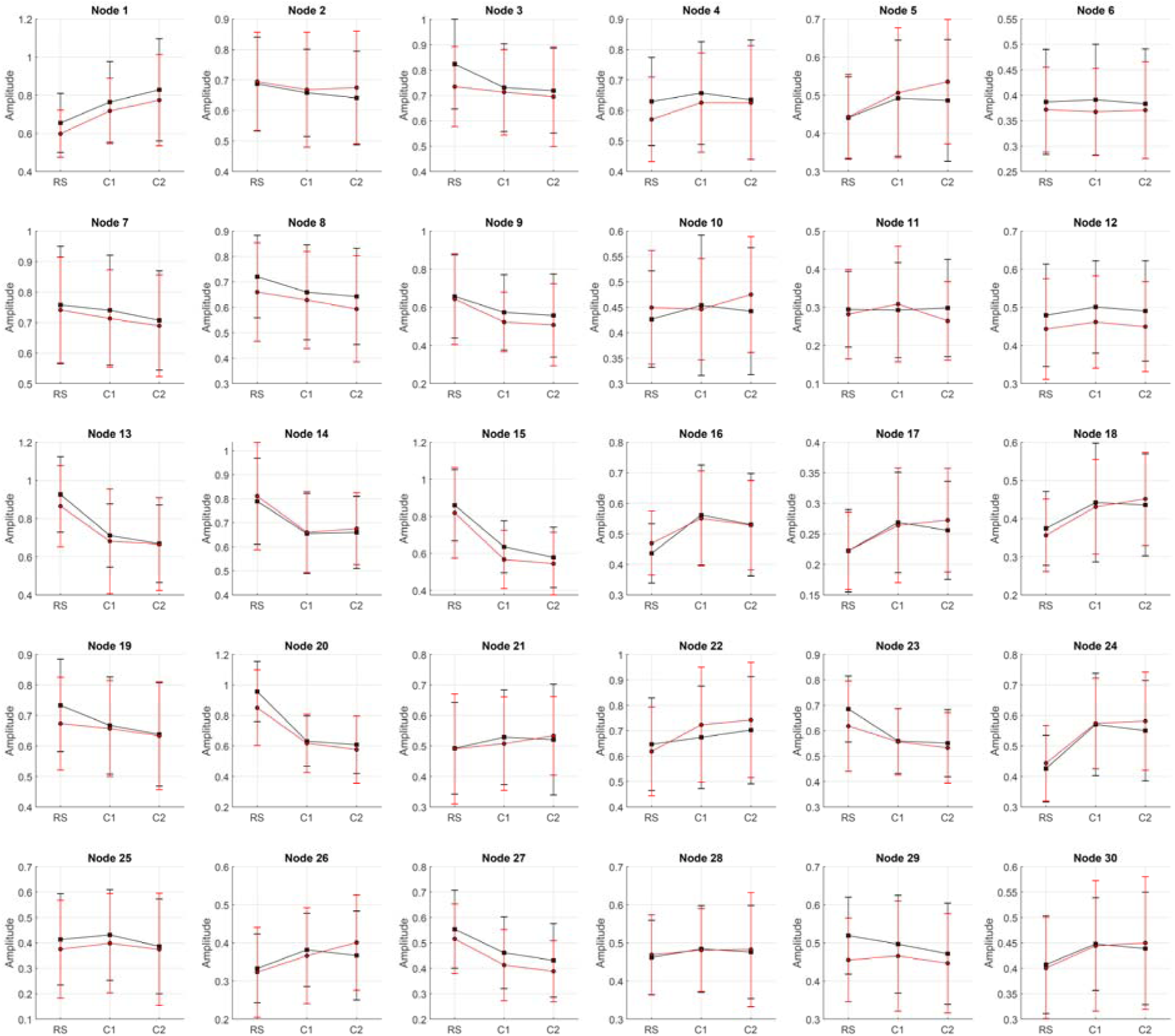
Visualization of nodewise SDSA for the 30 non-noise components for stroke patients (red lines) and healthy controls (black lines) during resting state and two load levels of the continuous tracking task.

## Discussion

Identifying sensitive imaging markers for early evaluation of severity and prognosis is important for improving patient stratification and personalized approaches in stroke care. In an attempt to classify patients with sub-acute strokes from healthy controls, we used fMRI-based brain network approaches to estimate indices of brain functional connectivity during an unconstrained resting state, and during two load levels of a multiple object tracking task. In line with our previous studies (Dørum et al., 2017), the classification analysis successfully distinguished between resting state and attentive tracking, with accuracies approaching 100% across groups. Despite the high sensitivity to experimental condition, and contrary to our initial hypothesis, the algorithm was not able to robustly distinguish between patients and healthy controls. Based on a notion of effort-dependent aberrations in brain functional organization, we had anticipated that increasing cognitive demand would increase the ability to discriminate between groups, however, the results did not support this hypothesis as increasing load levels had negligible effect on classification accuracy.

The results obtained using machine learning based classification were largely supported by the univariate edgewise analyses. Edge-level findings revealed strong main effects of condition on a range of edges, only one edge showing significant main effects of group, and no significant interactions between group and condition. The strong effects of experimental condition were corroborated in our node-wise analysis, showing robust modulation of task condition on SDSA in 19 of the 30 nodes, but no significant group differences.

In our previous study (Dørum et al., 2017), multivariate classification using the full set of FC indices yielded robust discrimination between groups of younger and older healthy adults as well as between states of rest and task engagement, thus providing a sensitive framework in which to explore age-related changes in neurobiology and their interactions with cognitive states. Correspondingly - employing the same classification analysis - we expected to find robust discrimination between a group of stroke sufferers and healthy adults. Whereas results in this study indicate high accuracy when classifying between resting state and two levels of attentional demand; group discrimination performed at chance-level. Machine learning and pattern recognition algorithms enable the capture of multivariate associations beyond traditional univariate analyses and are thus sensitive to differences in spatially distributed patterns of FC, which may serve as noninvasive biomarkers for disease. This methodological approach is congruent with the contemporary view of the brain as an integrative network, and has proven sensitive to distinguish group differences in multiple disease states (Arbabshirani et al., 2013; Craddock et al., 2009; Plitt et al., 2015).

Considering the dependency of FC on cognitive context, we hypothesized that a constrained cognitive task paradigm would comprise a more sensitive context for the study of brain network alterations after a cerebral stroke. In our previous study (Dørum et al., 2017), we found this paradigm to increase classification accuracy between a group of older and a group of younger healthy adults, and thus we expected to find a similar trend in classifying between a group of stroke patients and a matched cohort of healthy adults. Results did not support this hypothesis, however, as group classification accuracy remained largely unchanged from resting-state to the two load levels of the attention task.

It is possible that the poor group discriminability can be partly explained by the relative high functioning of the stroke sample. The inherent demands of the study biased patient selection towards less clinically severe strokes – reflected in an average NIHSS score below 1. Further, although healthy controls on average performed better than the stroke group in the cognitive assessment tests as well as during both load levels of the MOT-task, as indexed by response accuracy, these group differences were not found to be statistically significant. Thus, neuropsychological data corroborated the classification analyses with results indicating that clinically mild strokes may result in minimal behavioral and neural network-level effects beyond the area of the lesion. Another, not mutually exclusive aspect of the patient sample is a high level of premorbid functioning. It is possible that patients with higher cognitive capacity are more motivated and able to participate compared with the general stroke population, and that mechanisms related to their premorbid functioning are involved in explaining the apparent null effect. This hypothesis needs to be tested in future studies including lower functioning patients.

Multivariate analysis revealed chance-level discrimination between the stroke group and the healthy controls. Univariate repeated measures ANOVAs probing group differences revealed a significant main effect of group in an edge connecting a temporal and a sensorimotor node where connectivity strength was stronger in the stroke patients than the healthy controls.

Whereas group differences were subtle, a strong network response was observed across groups when participants were engaged in state of effortful attention. The repeated measures ANOVA identified significant main effects of task condition in several edges. The DAN was the network mostly implicated, showing a task-related reduction in connectivity with nodes in the somatosensory, supramarginal and fusiform cortices, as well as a task-related increase in connectivity with a left-lateralized frontoparietal node. Indicating reduced temporal coherence between task-relevant and task-irrelevant networks and increase coherence within task-relevant networks when participants were in a state of effortful attention. Our analyses revealed no significant interactions between group and task condition, suggesting similar task modulation in FC in the two groups.

Using a similarly heterogenous sample of stroke patients 1-2 weeks post-stroke incident, Baldassare and colleagues (2014) found an association between severity of post-stroke deficits and reduction in interhemispheric FC between the DAN and sensorimotor networks as well as an increase in FC between the usually anticorrelated DAN and DMN. Negative correlation between the internally oriented DMN and the externally focused DAN suggests a competitive relationship between these networks, and reduced anticorrelation or node differentiation has been observed in advancing age (Dørum et al., 2016), Alzheimer’s Disease (Weiler et al., 2017) and Parkinson’s Disease (Baggio et al., 2015), and might thus reflect neural network degeneration.

Brain signal variability facilitates transition between network configurations and reflects neural network adaptability and efficiency to respond to a greater range of stimuli. Lifespan developmental trajectories conform to an inverted U-shaped curve with less variability in the extremes of age and higher variability during adulthood (Garrett et al., 2010; McIntosh et al., 2008) similar to the quadratic trajectories observed for a range of cerebral health measures such as white matter properties (Westlye et al., 2010; Yeatman et al., 2014) and network modularity (Zuo et al., 2017). Studies have reported a positive association between BOLD signal variability and superior, as well as more consistent performance on a range of cognitive tasks (Garrett et al., 2011, 2012). Thus, we hypothesized higher signal variability for healthy adults compared to stroke patients, reflecting healthier neural dynamics. However, commensurate with edge-level results, our findings revealed no significant group differences in nodal signal variability, further corroborating the notion that clinically mild strokes yield minimal whole-brain neuronal effects.

Whereas group differences were indiscernible, task effects were substantial, reflected in 19 out of 30 nodes showing significant task modulation on node SDSA. Strongest effects were observed in task-positive frontoparietal networks and the task-negative DMN, as well as the cerebellum. The effects of task were consistent with findings in our previous study, where we observed both task-induced increases and decreases in signal variability for task-positive nodes, and a uniform decrease in SDSA for the DMN.

The present study should be interpreted with certain considerations in mind. The lack of discernable FC alterations between stroke patients and healthy controls might in part be explained by the patient recruitment procedure, specifically the lengthy and demanding task-fMRI session requisite for study participation. Considerable demand was placed on the patients having intact cognitive, motor and visual functions shortly following the stroke incident. As a result, the final patient selection did not reflect a representative cross-section of stroke patients admitted to stroke wards, but rather patients with clinically mild strokes which were exclusively of ischemic etiology, as ischemic strokes are both more common and less debilitating than hemorrhagic strokes (Andersen et al., 2009). A possible methodological explanation for the absence of significant group differences is the dimensionality of the networks derived from the ICA. We estimated networks at a model order of 40 components. The ability to detect group differences in FC may vary as a function of ICA model order (Abou Elseoud et al., 2011), and future studies in larger samples may be able to test the sensitivity to group differences across a range of model orders.

In conclusion, the main findings in this study demonstrate that FC patterns between a group of sub-acute stroke patients and a group of healthy controls were indiscernible using multivariate machine learning classification. While we observed high classification accuracy between data obtained during an unconstrained resting state and data obtained during a constrained attentive tracking task, this increase in cognitive demand did not yield an increase in group classification accuracy. Complimentary node-level analyses corroborated the edge-level findings converging on minimal whole-brain neuronal effects of clinically mild ischemic strokes.

